# A Complex Evolutionary History for the Disease Susceptibility *CDHR3* Locus

**DOI:** 10.1101/186031

**Authors:** Mary B. O’Neill, Guillaume Laval, João C. Teixeira, Ann C. Palmenberg, Caitlin S. Pepperell

## Abstract

Selective pressures imposed by pathogens have varied among human populations throughout their evolution, leading to marked inter-population differences at some genes mediating susceptibility to infectious and immune-related diseases. A common polymorphism resulting in a C_529_ versus T_529_ change in the Cadherin-Related Family Member 3 (*CDHR3*) receptor is associated with rhinovirus-C (RV-C) susceptibility and severe childhood asthma. Given the morbidity and mortality associated with RV-C dependent respiratory infections and asthma, we hypothesized that the protective variant has been under selection in the human population. Supporting this idea, a recent cross-species outbreak of RV-C among chimpanzees in Uganda, which carry the ancestral ‘risk’ allele at this position, resulted in a mortality rate of 8.9%. Using publicly available genomic data, we sought to determine the evolutionary history and role of selection acting on this infectious disease susceptibility locus. The protective variant is the derived allele and is found at high frequency worldwide, with the lowest relative frequency in African populations and highest in East Asian populations. There is minimal population structure among haplotypes, and we detect genomic signatures consistent with a rapid increase in frequency of the protective allele across all human populations. However, given strong evidence that the protective allele arose in anatomically modern humans prior to their migrations out of Africa and that the allele has not fixed in any population, the patterns observed here are not consistent with a classical selective sweep. We hypothesize that patterns may indicate frequency-dependent selection worldwide. Irrespective of the mode of selection, our analyses show the derived allele has been subject to selection in recent human evolution.

## Introduction

There is accumulating evidence to suggest that immunity-related genes are preferential targets of natural selection (reviewed in ^1–5^), supporting the notion that infectious diseases have been important selective forces on human populations^6^. However, for most candidate loci, the mechanisms and phenotypic effects underlying the observed signatures of selection remain elusive.

Human rhinoviruses (RV) are found worldwide and are the predominant cause of the common cold. While many RV infections cause only minor illness, type C strains of the virus are associated with higher virulence. Unlike RV-A and RV-B, RV-C utilizes the Cadherin Related Family Member 3 (CHDR3) receptor to gain entry into host cells. A missense variant in *CDHR3* (rs6967330, C529T) results in a 10-fold difference in virus binding and replication *in vitro* ^7^, and experimental evidence points to varied cell surface expression between the two forms of the receptor as a likely mechanism^8^. Adding clinical support that CDHR3 functions as an RV-C receptor, the ancestral allele at rs6967330 has been found to be associated with an increased risk of respiratory tract illness by RV-C in two birth cohorts^9^. The allele is also associated with a severe form of childhood asthma, in accordance with the known role of RV, particularly RV-C, in triggering asthma exacerbations^10^. Taken together, these studies suggest that host genetics mediate differing susceptibility to RV-C infection and asthma by affecting interactions between the virus and its receptor (e.g. whether or not the protein is expressed on the cell surface and is thus visible to the virus).

A recent outbreak of lethal respiratory illness among wild chimpanzees was attributed to human RV-C crossing species boundaries. All members of the infected chimpanzee population were invariant for the ancestral variant at the homologous position corresponding to the human rs6967330 variant. The outbreak resulted in a staggering 8.9% mortality rate ^11^, and suggests that RV-C infection can result in a high mortality rate in a susceptible population. As respiratory infections, particularly prior to the availability of modern medical interventions, represent significant threats to human health^12–15^, *CDHR3*, and in particular rs6967330, represent a promising candidate target of natural selection. In the present study, we investigated the evolutionary history of this locus for which there already exists strong experimental and clinical data linking genotype with phenotypes that appear to modulate disease susceptibility.

## Results & Discussion

### Long-term evolution of the *CDHR3* locus

Examination of the locus in an alignment of 100 vertebrate genomes^16,17^ revealed that the *CDHR3* locus is highly conserved, with homologs present in 85 species (Figure 1). Tyrosine is the ancestrally encoded amino acid at the homologous position 529 in the human protein sequence. There appears to have been a mutation leading to an A529T amino acid change introduced in a common ancestor of opossums, Tasmanian devils, and wallabies that results in the encoding of a phenylalanine at the homologous position. Numerous vertebrates (elephant, horse, rabbit, pika, naked mole-rat, rat, mouse, golden hamster, Chinese hamster, prairie vole, and lesser Egyptian jerboa) spread throughout the species tree also encode a different amino acid (histidine) at this position. Whether these mutations are fixed in each of these species requires further sequencing, and the effect of these nonsynonymous substitutions on protein expression and function remain to be explored.

**Figure 1.**
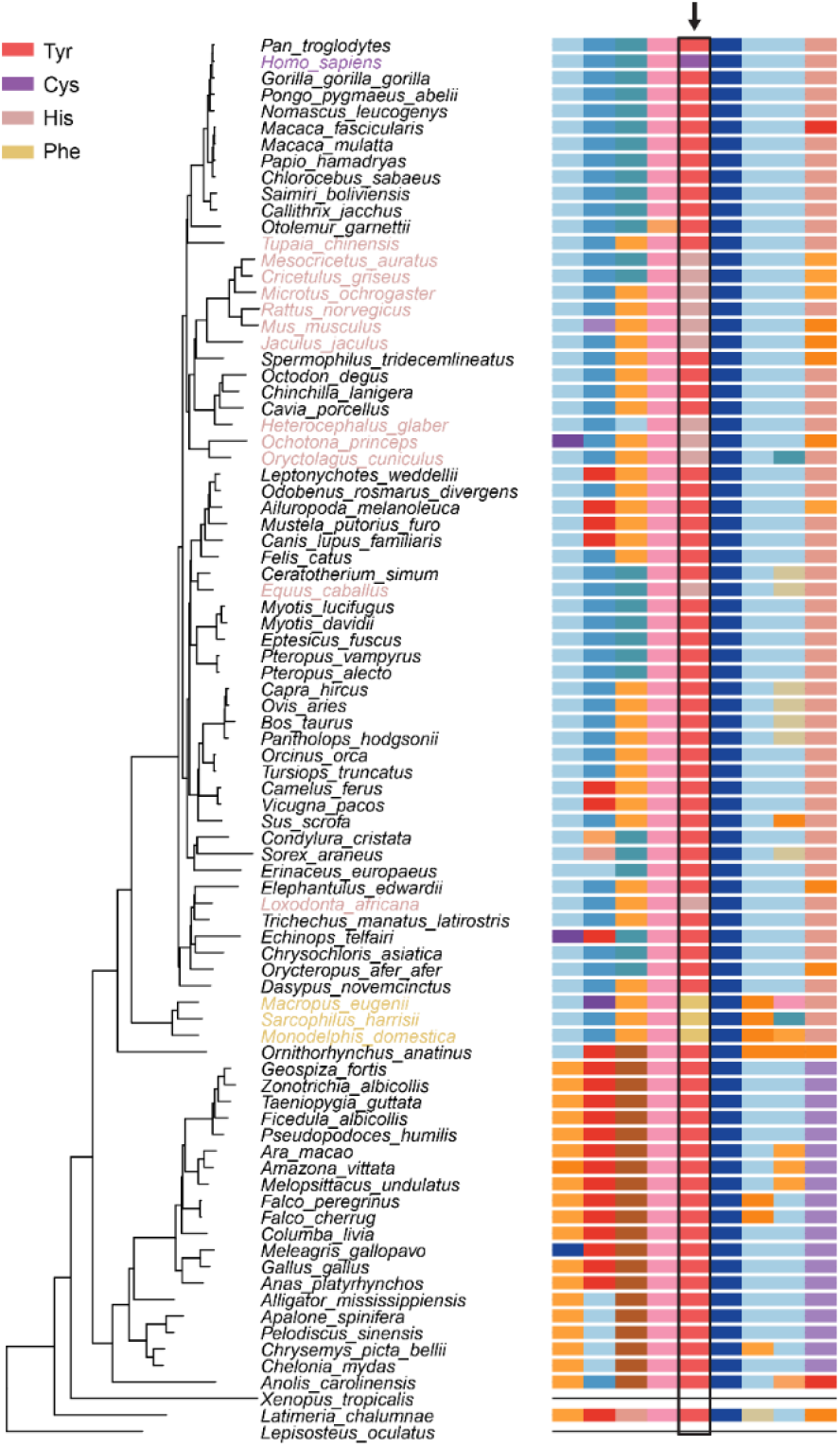
CDHR3 protein. (Left) Species tree for the multi sequence alignment of 85 species in the UCSC multiz alignment. (Right) Multi sequence protein alignment surrounding position 529 of the human CDHR3 protein. Residue 529 is outlined in black and designated with an arrow. Humans carry the Tyr and Cys alleles. Data obtained from the UCSC Genome Browser^16,17^.

Providing further evidence that the locus has been evolutionary constrained, rs6967330 has a positive Genomic Evolutionary Rate Profiling (GERP) score of 4.39, signifying a deficit of substitutions compared to neutral expectations. Values of the statistic calculated for bases within *CDHR3* range from −12.2 to 6.08 (mean = −0.15, sd = 2.59) and from −12.3 to 6.17 chromosome-wide (mean =-0.09, sd = 1.91)^17,18^. The apparent conservation of this locus suggests it is important for the function of CDHR3 and thus makes it an interesting candidate of natural selection.

### Population diversity patterns of the *CDHR3* locus

Excluding *Homo sapiens*, sequencing data from the remaining extant species comprising all hominids (great apes) are invariant at the position corresponding to rs6967330^19^. Genotyping of an additional 41 chimpanzees whose community experienced a severe respiratory outbreak of RV-C in 2013 revealed that all individuals were homozygous for the ancestral allele^11^. In examining hominin genomes, we find that Neanderthals and Denisovans carry the ancestral allele, while ancient specimens of anatomically modern humans carry both the ancestral and derived alleles. In low coverage sequencing data of *H. sapiens* estimated to have lived between 5000-8000 years ago (using sequencing reads from 45 aDNA samples for which we felt confident making diploid genotype calls) we estimate the allele frequencies at rs6967330 to be 34.4% A (ancestral) and 65.5% G (derived) (Table S1)^20^. Higher coverage aDNA extracted from a 7,000 year old skeleton found in Germany and an 8,000 year old skeleton from the Loschbour rock shelter in Luxembourg reveals heterozygotes at the locus^21^. Finally, a 45,000 year old early H. *sapiens* from western Siberia is a homozygote for the derived allele^22^. Collectively, these data suggest that the derived allele likely arose in the evolutionary branch leading to anatomically modern humans, although population data for Neanderthals and Denisovans remains scarce.

The locus represents a shared polymorphism in contemporary worldwide populations. Based on whole genome sequence data for 2504 individuals from 26 different populations^23^, we find that the derived (‘protective’) G allele is, on average, very common in East Asian populations (EAS, 93.0%), followed by admixed American populations (AMR, 85.7%), South Asian populations (SAS, 80.0%), and European populations (EUR, 79.2%). It is least common in African populations (AFR, 73.5%), albeit also at high frequency. At the individual population level, allele frequencies of the derived G allele range from 68.8% (“Mende in Sierra Leone”, MSL) to 95.3% (“Peruvians from Lima, Peru”, PEL) (Figure 2).

**Figure 2.**
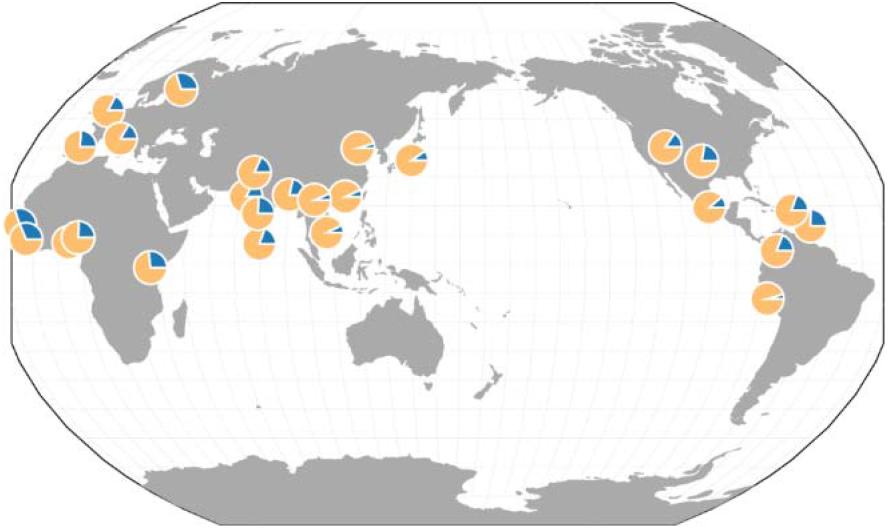
Global distribution of allele frequencies at rs6967330. Pie charts represent the relative allele frequencies of the ancestral A allele (blue) and the derived G allele (yellow) in each of the 26 populations of the 1000 Genomes Project^23^. This image was generated through the *Geography of Genetic Variants* browser^52^.

We next investigated linkage disequilibrium (LD) patterns around *CDHR3* with Haploview^24^. A tight LD block was identified on chromosome 7 from 105,657,078-105,659,873, with only moderate LD decay extending up to 105,680,022. Considering only biallelic SNPs with a minor allele frequency (MAF) > 0.01, haplotypes within these blocks were identified with the Pegas package in R^25,26^. Relationships among haplotypes occurring at >1% were inferred using network analysis (Figure 3). Eight haplotypes at frequency ≥ 1% were identified from the region of high LD, with the majority of individuals in all populations carrying the same haplotype with the derived G allele (*n* = 3848); two less frequent haplotypes carrying the derived allele are found in various geographic regions. The ancestral allele is found in the remaining five haplotypes, which vary in their distributions across regions. Twenty-eight haplotypes were found using the same criteria of a MAF > 0.01 and haplotype frequency of > 1% in the larger genomic block containing moderate LD. Phylogenetic reconstruction of the resulting haplotypes in this region resulted in a clear separation of those carrying the ancestral and derived alleles. Unexpectedly, Chimp, Neanderthal and Denisovan haplotypes are closer to *H. sapiens* A-haplotypes suggesting that natural selection shaped the haplotype pattern (Figure S1).

**Figure 3.**
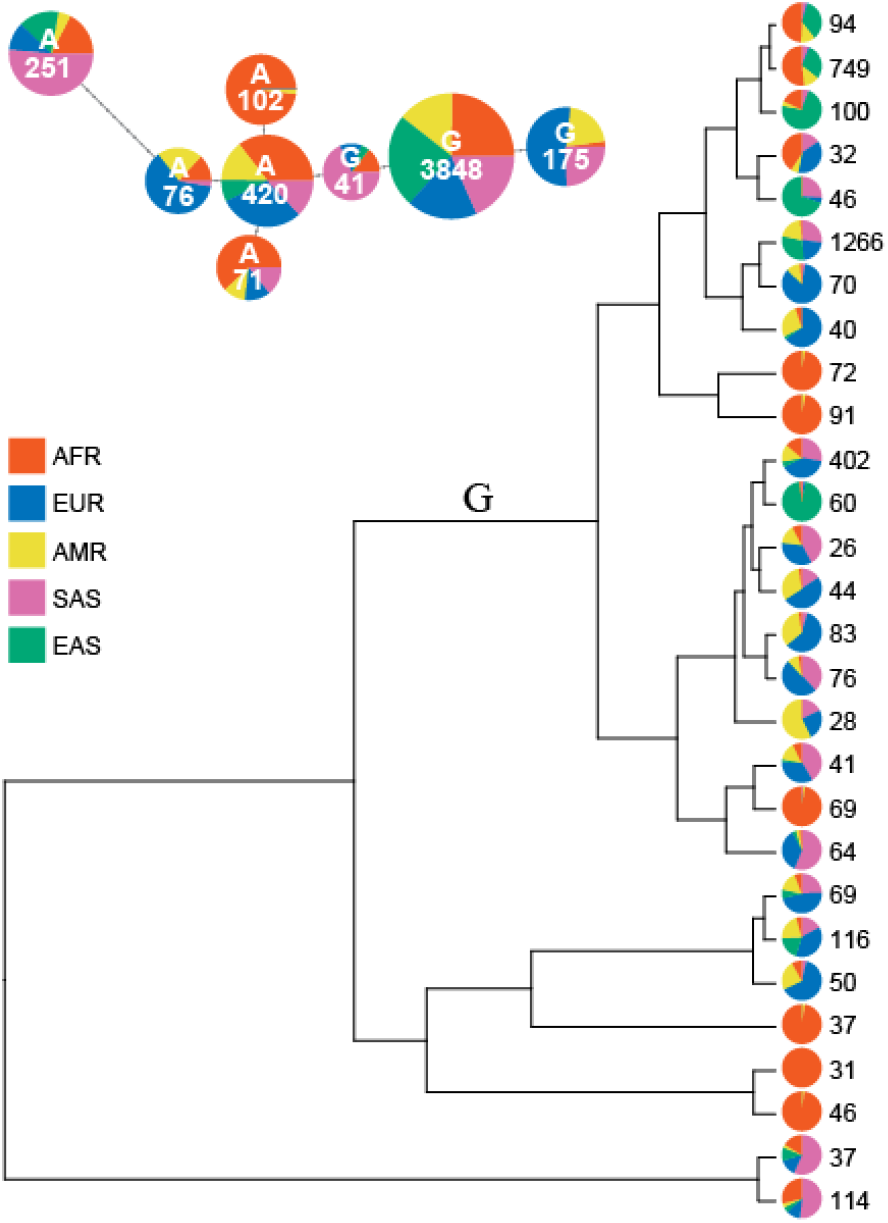
Haplotype structure of anatomically modern humans from the 1000 Genomes Project. (Top) Haplotype network from chromosome 7 between 105,657,078 and 105,659,873. Haplotype network was constructed from phased genome sequences of 2504 individuals with variation at 16 sites in the 2,795bp region. Only haplotypes occurring at > 1% are shown. Colors reflect super population designation of individuals. (Bottom) Unrooted tree from chromosome 7 between 105,657,078 and 105,680,022. Haplotypes of the same 2504 individuals were derived for a larger haplotype block. Again, only haplotypes occurring at > 1% are shown and colors reflect super population designation of individuals. The tree is based on 83 SNPs (68 informative) from the 22,945bp region.

### World-wide Selection at the locus

Given the morbidity and mortality associated with viral respiratory infections^13,15^ and severe childhood asthma^12,14^ particularly prior to the availability of modern medical interventions, we hypothesized that the frequency of the protective G allele (resulting in a Cys_529_) at rs6967330 might have increased more rapidly than under neutrality. We performed genome-wide scans for selection in the 26 populations of the 1000 Genomes Project^23^. To do so, we used the integrated haplotype score (iHS)^27^ and the number of segregating sites by length (*nS_L_*)^28^, which are designed to capture rapid increases in frequency of selected variants and have been applied in the context of detecting recent positive selection ^27,29,30^. We calculated these neutrality statistics in each population independently and normalized by frequency-bin for all loci with a DAF ≥ 0.2 (Figure S2). Fourteen populations presented extreme values of iHS and *nS_L_* (≥ 95^th^ percentile of the distribution in the population) at rs6967330, with 2 and 5 populations falling in the 99^th^ percentile for the two statistics, respectively. *CDHR3* lies in a region of high recombination (1.8cM/Mb) and rs6967330 resides in a male specific recombination hotspot^17,31^. This may explain the slightly different patterns observed in *nS_L_* and iHS statistics, as an excess of extreme values of iHS have been observed at regions of low recombination^28^.

To investigate whether selection has acted on all populations simultaneously, we implemented a multi-population (MP) combined statistic approach for iHS and *nS_L_*. For each SNP in each population, we combined the percent ranks of iHS and *nS_L_* using Fisher’s method, and defined the MP-iHS and MP-*nS_L_* as the resulting *χ*^2^ value. We obtained a MP-iHS score of 156 and a MP-*nS_L_* score of 157 for rs6967330. Both scores are in the 99^th^ percentile, regardless of whether we include only segregating sites in 2 or more populations or limit it to those SNPs segregating in all 26 populations examined. This suggests that the derived allele at rs6967330 is advantageous and that non-neutral processes have acted on this locus across global populations.

### A complex evolutionary scenario

We originally hypothesized that selection pressures imposed by RV-C infection may have resulted in a classic selective sweep at the locus; however, our results here are not consistent with such a model. Preliminary dating estimates of RV-C point to a recent origin in the last few thousand years^8^. We infer that the protective (derived) allele was already present in ancestral populations at intermediate frequencies during this time frame (65.5% in aDNA dated 5,000-8,000 years ago). Haplotype-based statistics such as iHS and *nS_L_* have low power to detect selection occurring on standing variation with intermediate frequency^32^. This suggests that patterns of variation observed at rs6967330 are unlikely to have resulted from selection imposed by RV-C itself; our findings suggest instead that an alternative selective agent(s) may have been/be acting on the *CDHR3* locus prior to emergence of RV-C. These findings well parallel those observed at the chemokine receptor gene-5 (*CCR-5*), where an unknown historical selective pressure maintained a deletion in this gene that attenuates infectivity and disease progression of HIV (reviewed in ^33^), because that mutation clearly predates the emergence of AIDS.

### Frequency-dependent Selection

Several lines of evidence point to the derived allele at rs6967330 arising before the out-of-Africa dispersal, including a lack of haplotype structure (Figure 3) and the presence of the allele at homozygous state in aDNA from anatomically modern humans dating back to ~45,000 year old^20–22^. While we find signatures of selection at the locus, the protective variant has not fixed in any of the contemporary populations examined (Figure 2), contrary to expectations for a strongly positively selected variant that arose prior to human migrations out of Africa^34^. We therefore wondered if balancing selection could explain observed patterns of variation at the locus, as balancing selection can maintain functional diversity over long periods of time through frequency-dependent selection, heterozygote advantage, pleiotropy, and fluctuating selection^35^. Until recently, evidence for balancing selection in the human genome was limited to a few classical cases such as the heterozygous advantage conferred by the *HbS* sickle cell mutation against malaria^36,37^, genes of the major histocompatibility complex/human leukocyte antigen complex^38^, and the ABO blood group^39^. Balancing selection, however, has recently been recognized as more prevalent than previously thought, particularly in shaping human immune system phenotypes^35,40,41^. To test whether the rs6967330 locus has been under long-term balancing selection (i.e. selection occurring over at least hundreds of thousands of generations ^38,42,43^), we examined the distribution of *β*, a recently developed summary statistic designed to detect clusters of alleles at similar frequencies, in 1000 Genomes Project data^44^. *β* was similar across all populations around the rs6967330 locus, and values were not indicative of long-term balancing selection (Figure S3).

Balancing selection operating over shorter timescales clearly also plays a role in shaping human diversity (e.g. *HbS* sickle cell mutation^36,37^). Such a short-term balancing selection scenario could explain the signals we detected with haplotype-based statistics, as genomic signatures of short-term balancing selection are predicted to be indistinguishable from incomplete sweeps of positive selection ^27,41^. In both of these scenarios, the selected allele rapidly increases in frequency, with the allele eventually fixing under positive selection or oscillating around a steady state in the case of balancing selection. As a reminder, haplotype-based selection methods suggest that rs6967330 rose in frequency faster than under neutrality, both at the global and the individual population level. Furthermore, population differentiation as measured with *F_ST_* at this locus is low relative to other SNPs with the same frequency (Figure S4), a finding that is also consistent with short-term balancing selection across multiple populations^35,38,41^. Based on our observation that the derived variant at rs6967330 was present at high frequencies in ancient human specimens and in the homozygous state in the case of a 45,000-year-old fossil, we posit that this allele may have started to increase in frequency in ancestral populations before reaching its current equilibrium frequency independently in worldwide populations. The derived allele ranges in frequency from 68.8-95.3% in modern human populations, which is compatible with an equilibrium frequency being high. It is possible that as the allele frequency of the derived mutation reaches high frequencies in a population, circulation of the viruses exploiting CDHR3 (past and present) are affected, with attendant impacts on the selective pressures they impose on human populations. Such a frequency-dependent selection scenario seems more plausible than that of a heterozygote advantage given that in Danish children with severe asthma, having even one copy of the risk variant was associated with increased risk of exacerbation and hospitalization^10^. If there were another virus that exploited this receptor in past human populations, we might assume that heterozygotes would similarly not be conferred protection.

In conclusion, our analyses combined *a priori* knowledge of a genetic variant underlying variable susceptibility to RV-C infections^7,9,11^ with population genetic analyses of whole genome sequence data to investigate the evolutionary history of the locus in *CDHR3*. The conservation of the protein, combined with its complex evolutionary history, exemplifies the biological importance of *CDHR3*, which may or may not be ultimately relevant to its function as the cellular surface receptor for RV-C. We detected a worldwide signature of selection at this locus, but also found that patterns of variation do not conform to the classic selective sweep model. Instead, we posit the possibility of frequency-dependent balancing selection operating at this locus which warrants more investigation into genotypic and biochemical effects of the variant (e.g. whether there is any phenotypic benefit to being heterozygous, trade-offs of different genotypes, etc.). Irrespective of the mode of selection, our analyses show that the derived allele has been advantageous and selected for in recent human evolution.

## Methods

### Datasets

#### 1000 Genomes Project

Individual level phased sequencing data from the 1000 Genomes Project Phase 3 dataset were downloaded from ftp://ftp.1000genomes.ebi.ac.uk/vol1/ftp/release/20130502/^23^.

#### Ancient H. sapiens

BAM files for each of the 230 individuals included in the study by Mathieson *et. Al.*^20^ were downloaded and converted to MPILEUP format using samtools^45^. Low coverage sequencing data at asthma susceptibility loci were manually inspected for each individuals for which at least one read had mapped to the site. Manual diploid genotyping calls were made for aDNA samples for which we felt confident making diploid genotype calls (Table S1).

#### Neandertal and Denisovan

Geontypes for the Vindija, Altai, and Denisovan genomes^46^ generated using snpAD, an ancient DNA damage-aware genotyper, were downloaded from http://cdna.eva.mpg.de/neandertal/Vindija/VCF/.

#### Great Apes

Genotypes of primates sequences were obtained from https://eichlerlab.gs.washington.edu/greatape/data/ and converted to the corresponding human regions with the LiftOver software^17^.

### Haplotype Networks

Indels and multi-allelic sites were filtered out with bcftools^45^. Variants having a Hardy-Weinberg equilibrium exact test p-value below 1×10^−5^ as calculated using the – hwe midp function in PLINK1.9^47^ in any of the 26 populations were removed from all populations. The core haplotype surrounding rs6967330 was identified using biallelic markers within 100kb of rs6967330 in Haploview^24^ from all 26 populations in the 1000 Genomes Phase 3 release. A large haplotype block was defined on Chromosome 7 from 105,657,078 to 105,680,022, and a smaller haplotype of Chromosome 7 from 105,657,078 to 105,659,873. Haplotypes within the defined haplotype blocks were extracted from biallelic markers with a minor allele frequency (MAF) > 0.01 with the Pegas package^25,26^. Haplotypes occurring at >1% (at least 26 individuals) in the total 1000 Genomes Project dataset were constructed into networks. Genotypes from two high quality Neanderthal genomes and a Denisovan genome were similarly extracted and used in network analyses.

### Haplotype-Based Selection Scans

We computed iHS and *nS_L_* using the method implemented in Selink, a software to detect selection using whole-genome datasets (https://github.com/h-e-g/selink). As iHS and *nS_L_* are sensitive to the inferred ancestral/derived state of an allele, we computed these statistics only when the derived state was determined unambiguously^32^. Results were normalization by derived allele frequency (DAF) bins (from 0 to 1, increments of 0.025)^27,32^. We also minimized the false-positive discovery by excluding SNPs with a DAF below 0.2, as the power to detect positive selection has been shown to be limited at such low frequencies^27,32^. For each statistic, we considered the percent rank at rs6967330 relative to the genome-wide distribution in each population.

In order to test selection shared among populations, we combined iHS and *nS_L_* into a single multi-population (MP) combined selection score, MP-iHS and MP-*nS_L_*. The rationale behind these composite approaches^48–50^ is that neutrality statistics, though expected to be correlated among populations under neutrality, are more strongly correlated for positively selected variants than for neutral variants. Indeed, under global positive selection, iHS and *nS_L_* tend to become negative in all populations while false positives will only be negative in a few populations. Consequently, candidates genuinely selected in several populations should harbor extreme values for MP-iHS and MP-*nS_L_*. For each SNP and each population, we determined the empirical *p*-value of iHS and *nS_L_*, i.e. the rank of each statistic in the genome-wide distribution divided by the number of SNPs. We then used the combine_pvalues function implementation of Fisher’s method^51^ of the SciPy python package to compute MP-iHS and MP-nS_L_ for every SNP (passing the criteria above to compute iHS and *nS_L_*) found in at least two populations.

### *β* Test for Long-Term Balancing Selection

*β* scores for each population in the 1000 Genomes Project were obtained from https://github.com/ksiewert/BetaScan.

### Population Differentiation

Global *F_ST_* was calculated with Selink (https://github.com/h-e-g/selink).

## Acknowledgements

We wish to thank Andrew Kitchen for his advice on phylogenetic analyses and Lluis Quintana-Murci for feedback on the manuscript.

## Author Contributions

MBO, CSP, and ACP conceived the project. MBO, GL, and JCT performed the analyses. MBO drafted the paper and made the figures. All authors provided critical feedback, reviewed, and edited the manuscript.

## References

1. Barreiro, L. B. & Quintana-Murci, L. From evolutionary genetics to human immunology: how selection shapes host defence genes. Nat Rev Genet 11, 17–30 (2010).

2. Siddle, K. J. & Quintana-Murci, L. The Red Queen’s long race: human adaptation to pathogen pressure. Current Opinion in Genetics & Development 29, 31–38 (2014).

3. Karlsson, E. K., Kwiatkowski, D. P. & Sabeti, P. C. Natural selection and infectious disease in human populations. Nat Rev Genet 15, 379–393 (2014).

4. Fumagalli, M. & Sironi, M. Human genome variability, natural selection and infectious diseases. Current Opinion in Immunology 30, 9–16 (2014).

5. Quach, H. & Quintana-Murci, L. Living in an adaptive world: Genomic dissection of the genus Homo and its immune response. Journal of Experimental Medicine 214, 877–894 (2017).

6. Fumagalli, M. et al. Signatures of Environmental Genetic Adaptation Pinpoint Pathogens as the Main Selective Pressure through Human Evolution. PLOS Genetics 7, e1002355 (2011).

7. Bochkov, Y. A. et al. Cadherin-related family member 3, a childhood asthma susceptibility gene product, mediates rhinovirus C binding and replication. PNAS 112, 5485–5490 (2015).

8. Palmenberg, A. C. Rhinovirus C, Asthma, and Cell Surface Expression of Virus Receptor, CDHR3. J. Virol. JVI. 00072–17 (2017). doi:10.1128/JVI.00072-17

9. Bønnelykke, K. et al. Cadherin-related Family Member 3 Genetics and Rhinovirus C Respiratory Illnesses. Am J Respir Crit Care Med 197, 589–594 (2017).

10. Bønnelykke, K. et al. A genome-wide association study identifies CDHR3 as a susceptibility locus for early childhood asthma with severe exacerbations. Nat Genet 46, 51–55 (2014).

11. Scully, E. J. et al. Lethal Respiratory Disease Associated with Human Rhinovirus C in Wild Chimpanzees, Uganda, 2013. Emerg Infect Dis 24, 267–274 (2018).

12. Gern, J. E. The ABCs of Rhinoviruses, Wheezing, and Asthma. J. Virol. 84, 7418–7426 (2010).

13. Bryce, J., Boschi-Pinto, C., Shibuya, K. & Black, R. E. WHO estimates of the causes of death in children. The Lancet 365, 1147–1152 (2005).

14. Busse, W. W., Lemanske, R. F. & Gern, J. E. The Role of Viral Respiratory Infections in Asthma and Asthma Exacerbations. Lancet 376, 826–834 (2010).

15. Ferkol, T. & Schraufnagel, D. The Global Burden of Respiratory Disease. Annals ATS 11, 404–406 (2014).

16. Karolchik, D. et al. The UCSC Table Browser data retrieval tool. Nucleic Acids Res. 32, D493–496 (2004).

17. Rosenbloom, K. R. et al. The UCSC Genome Browser database: 2015 update. Nucleic Acids Res 43, D670–D681 (2015).

18. Cooper, G. M. et al. Distribution and intensity of constraint in mammalian genomic sequence. Genome Res. 15, 901–913 (2005).

19. Prado-Martinez, J. et al. Great ape genetic diversity and population history. Nature 499, 471–475 (2013).

20. Mathieson, I. et al. Genome-wide patterns of selection in 230 ancient Eurasians. Nature advance online publication, (2015).

21. Lazaridis, I. et al. Ancient human genomes suggest three ancestral populations for present-day Europeans. Nature 513, 409–413 (2014).

22. Fu, Q. et al. The genome sequence of a 45,000-year-old modern human from western Siberia. Nature 514, 445–449 (2014).

23. The 1000 Genomes Project Consortium. A global reference for human genetic variation. Nature 526, 68–74 (2015).

24. Barrett, J. C., Fry, B., Maller, J. & Daly, M. J. Haploview: analysis and visualization of LD and haplotype maps. Bioinformatics 21, 263–265 (2005).

25. Paradis, E. pegas: an R package for population genetics with an integrated–modular approach. Bioinformatics 26, 419–420 (2010).

26. R Development Core Team. R: A Language and Environment for Statistical Computing. (R Foundation for Statistical Computing).

27. Voight, B. F., Kudaravalli, S., Wen, X. & Pritchard, J. K. A Map of Recent Positive Selection in the Human Genome. PLoS Biol 4, e72 (2006).

28. Ferrer-Admetlla, A., Liang, M., Korneliussen, T. & Nielsen, R. On detecting incomplete soft or hard selective sweeps using haplotype structure. Mol Biol Evol msu077 (2014). doi:10.1093/molbev/msu077

29. Sabeti, P. C. et al. Detecting recent positive selection in the human genome from haplotype structure. Nature 419, 832–837 (2002).

30. Ferrari, S. L., Ahn-Luong, L., Garnero, P., Humphries, S. E. & Greenspan, S. L. Two Promoter Polymorphisms Regulating Interleukin-6 Gene Expression Are Associated with Circulating Levels of C-Reactive Protein and Markers of Bone Resorption in Postmenopausal Women. J Clin Endocrinol Metab 88, 255–259 (2003).

31. Kong, A. et al. Fine-scale recombination rate differences between sexes, populations and individuals. Nature 467, 1099–1103 (2010).

32. Fagny, M. et al. Exploring the Occurrence of Classic Selective Sweeps in Humans Using Whole-Genome Sequencing Data Sets. Mol Biol Evol 31, 1850–1868 (2014).

33. Arenzana-Seisdedos, F. & Parmentier, M. Genetics of resistance to HIV infection: Role of co-receptors and co-receptor ligands. Seminars in Immunology 18, 387–403 (2006).

34. Stephan, W. Signatures of positive selection: from selective sweeps at individual loci to subtle allele frequency changes in polygenic adaptation. Molecular Ecology 25, 79–88 (2016).

35. Key, F. M., Teixeira, J. C., de Filippo, C. & Andrés, A. M. Advantageous diversity maintained by balancing selection in humans. Current Opinion in Genetics & Development 29, 45–51 (2014).

36. Allison, A. C. Protection Afforded by Sickle-cell Trait Against Subtertian Malarial Infection. Br Med J 1, 290–294 (1954).

37. Shriner, D. & Rotimi, C. N. Whole-Genome-Sequence-Based Haplotypes Reveal Single Origin of the Sickle Allele during the Holocene Wet Phase. The American Journal of Human Genetics 102, 547–556 (2018).

38. Leffler, E. M. et al. Multiple Instances of Ancient Balancing Selection Shared Between Humans and Chimpanzees. Science 339, 1578–1582 (2013).

39. Ségurel, L. et al. The ABO blood group is a trans-species polymorphism in primates. PNAS 109, 18493–18498 (2012).

40. Ferrer-Admetlla, A. et al. Balancing Selection Is the Main Force Shaping the Evolution of Innate Immunity Genes. The Journal of Immunology 181, 1315–1322 (2008).

41. Andrés, A. M. et al. Targets of Balancing Selection in the Human Genome. Mol Biol Evol 26, 2755–2764 (2009).

42. Wiuf, C., Zhao, K., Innan, H. & Nordborg, M. The Probability and Chromosomal Extent of *trans*-specific Polymorphism. Genetics 168, 2363–2372 (2004).

43. Teixeira, J. C. et al. Long-Term Balancing Selection in LAD1 Maintains a Missense Trans-Species Polymorphism in Humans, Chimpanzees, and Bonobos. Mol Biol Evol 32, 1186–1196 (2015).

44. Siewert, K. M. & Voight, B. F. Detecting Long-Term Balancing Selection Using Allele Frequency Correlation. Mol Biol Evol 34, 2996–3005 (2017).

45. Li, H. et al. The Sequence Alignment/Map format and SAMtools. Bioinformatics 25, 2078–2079 (2009).

46. Prüfer, K. et al. A high-coverage Neandertal genome from Vindija Cave in Croatia. Science eaao1887 (2017). doi:10.1126/science.aao1887

47. Purcell, S. et al. PLINK: a tool set for whole-genome association and population-based linkage analyses. Am J Hum Genet 81, 559–75 (2007).

48. Deschamps, M. et al. Genomic Signatures of Selective Pressures and Introgression from Archaic Hominins at Human Innate Immunity Genes. Am J Hum Genet 98, 5–21 (2016).

49. Grossman, S. R. et al. A Composite of Multiple Signals Distinguishes Causal Variants in Regions of Positive Selection. Science 327, 883–886 (2010).

50. Grossman, S. R. et al. Identifying Recent Adaptations in Large-Scale Genomic Data. Cell 152, 703–713 (2013).

51. Fisher, R. A. Statistical Methods for Research Workers. in Breakthroughs in Statistics 66–70 (Springer, New York, NY, 1992). doi:10.1007/978-1-4612-4380-9_6

52. Marcus, J. H. & Novembre, J. Visualizing the geography of genetic variants. Bioinformatics 33, 594–595 (2017).

